# Deep phenotyping of a healthy human HAO1 knockout informs therapeutic development for primary hyperoxaluria type 1

**DOI:** 10.1101/524256

**Authors:** Tracy L. McGregor, Karen A. Hunt, Paul Nioi, Dan Mason, Simina Ticau, Marissa Pelosi, Perry R. Loken, Sarah Finer, Christopher J Griffiths, Daniel G MacArthur, Richard C Trembath, Devin Oglesbee, John C. Lieske, John Wright, David V. Erbe, David A. van Heel

## Abstract

Primary Hyperoxaluria Type 1 (PH1) is a rare autosomal recessive metabolic disorder of oxalate metabolism leading to kidney failure as well as multi-organ damage. Overproduction of oxalate occurs in the liver due to an inherited genetic defect in the enzyme alanine-glyoxylate aminotransferase (*AGXT*), causing pathology due to the insolubility of calcium oxalate crystals in body fluids. The main current therapy is dual liver-kidney transplant, which incurs high morbidity and has poor availability in some health systems where PH1 is more prevalent. One approach currently in active clinical investigation targets *HAO1* (hydroxyacid oxidase 1), encoding glycolate oxidase, to reduce substrate levels for oxalate production. To inform drug development, we sought individuals with reduced HAO1 function due to naturally occurring genetic variation.

Analysis of loss of function variants in 141,456 sequenced individuals suggested individuals with complete *HAO1* knockout would only be observed in 1 in 30 million outbred people. However in a large sequencing and health records program (Genes & Health), in populations with substantial autozygosity, we identified a healthy adult individual predicted to have complete knockout of *HAO1* due to an ultra rare homozygous frameshift variant (rs1186715161, ENSP00000368066.3:p.Leu333SerfsTer4). Primary care and hospital health records confirmed no apparently related clinical phenotype. At recall, urine and plasma oxalate levels were normal, however plasma glycolate levels (171 nmol/mL) were 12 times the upper limit of normal in healthy, reference individuals (mean+2sd=14 nmol/mL, n=67) while her urinary glycolate levels were 6 times the upper limit of normal. Comparison with preclinical and phase 1 clinical trial data of an RNAi therapeutic targeting *HAO1* (lumasiran) suggests the individual likely retains <2% residual glycolate oxidase activity.

These results provide important data to support the safety of *HAO1* inhibition as a potential chronic therapy for a devastating metabolic disease (PH1). We also suggest that the effect of glycolate oxidase suppression in any potential other roles in humans beyond glycolate oxidation do not lead to clinical phenotypes, at least in this specific individual. This demonstrates the value of studying the lifelong complete knockdown of a target protein in a living human to aid development of a potential therapeutic, both in de-risking the approach and providing potential hypotheses to optimize its development. Furthermore, therapy for PH1 is likely to be required lifelong, in contrast to data from chronicity studies in non-human species or relatively short-term therapeutic studies in people. Our approach demonstrates the potential for improved drug discovery through unlocking relevant evidence hiding in the diversity of human genetic variation.

## INTRODUCTION

Primary Hyperoxaluria Type 1 (PH1) is a rare autosomal recessive metabolic disorder of oxalate metabolism with prevalence of 1-3/1,000,000 in Europe, accounting for 1-2% of paediatric end stage kidney disease, but with higher prevalence in some Middle East and North African countries^1^. Overproduction of oxalate occurs in the liver due to an inherited genetic defect in the enzyme alanine-glyoxylate aminotransferase (encoded by *AGXT*). Pathology ensues because of the increased oxalate that is unable to be further metabolized and must be excreted renally. The high concentrations of oxalate in the urine causes precipitation of highly insoluble oxalate salts (e.g. calcium oxalate) leading to a phenotype which varies from kidney failure and life-threatening consequences in infancy to occasional stone formation in adulthood. The earliest symptoms among those affected are kidney, urinary tract and bladder stones, leading to progressive kidney involvement and chronic kidney disease for most individuals. The disease course can also lead to multi-organ damage from systemic oxalosis, affecting bones, eyes, blood vessels, heart, thyroid, skin, and other tissues. The main definitive therapy is a dual liver-kidney transplant with long-term immunosuppression, which is dependent on a suitable donor being found, carries substantial morbidity and mortality, and can be unavailable in developing country health systems for some of the populations that are most affected. Thus, substantial need exists for therapies to treat PH1 without requiring liver transplantation.

One approach currently in active clinical investigation targets *HAO1* (hydroxyacid oxidase 1) encoding glycolate oxidase to reduce substrate for oxalate production in the liver. Glycolate oxidase is co-localised in the peroxisome of hepatocytes and is directly upstream of the AGT enzyme that is defective in PH1. By decreasing glycolate oxidase activity, oxalate production is expected to decrease, with an associated increase in glycolate levels. Glycolate is expected to be freely excreted by the kidneys due to its high solubility. However, suppression of glycolate oxidase is a novel approach to therapy for a metabolic disease, and therefore poses potential (and unknown) risks in terms of its chronic inhibition in humans. For example, glycolate oxidase is also active on 2-hydroxy fatty acids and its manipulation may affect their homeostasis. Pre-clinical examination of mice, rats, and monkeys following RNAi therapeutic-mediated silencing of *HAO1* has been completed with no suggestive adverse signals from chronic *HAO1* silencing of >95%, with the expected substantial elevations in glycolate levels^2^.

A powerful approach to target de-risking has been the identification of healthy individuals who spontaneously lack activity for a given protein (human knockouts), with biallelic *PCSK9* mutations resulting in deficiency as a classic example, where low LDL cholesterol and protection from cardiovascular disease is seen^3^. Additional recent examples include *LPA* knockouts that also confer protection from cardiovascular disease^4^, complete APOC3 deficiency providing protection against myocardial infarction^5^, a human with a *PRDM9* knockout with an unexpected meiotic recombination pattern but a fertile and healthy phenotype^6^, ANGPTL3 deficiency associated with decreased levels of all three major lipid fractions and decreased risk of atherosclerotic cardiovascular disease^7^. Such reverse genetics - identifying individuals with extreme genotypes and then investigating their biology, provides an efficient alternative to decades of traditional forward genetics - aggregating phenotypes and then investigating their genetic basis. Such human knockouts are especially informative about human biology, and are also uniquely useful for drug target validation and development.

A challenge to the use of reverse genetics more broadly, of course, stems from the reality that in the general population such individuals with homozygous or compound heterozygous deficiency are expected to be extremely rare and their identification is compounded when the interest is in clinically silent phenotypes (e.g. to de-risk therapeutic targets). Despite these challenges, the key question for *HAO1* therapeutic targeting is the long term safety of glycolate oxidase knockdown in humans, including any effects of elevated glycolate levels. Here, additional evidence from humans is clearly invaluable as naturally occurring genetic variation in humans provides opportunities to study the mechanism and safety of germline gene knockdown across the full life course of an individual^6,8^.

We therefore searched for naturally occuring *HAO1* knockouts in humans. No complete knockout (homozygous predicted loss of function) variants in *HAO1* were observed in the online gnomAD v2.1 database of 141,456 whole exome/genome sequenced individuals^9,10^. A literature search revealed three case reports of potential HAO1 null individuals, all in children and all either complicated by other diseases/genetics and/or missing important information. Frishberg *et al.* reported an 8 year old boy born to consanguineous parents who was investigated for psychomotor delay, anisocoria and alacrima and diagnosed with triple-A-like syndrome due to homozygous predicted loss of function (pLoF) mutations in *GMPPA*^*11*^. During the course of his diagnostic evaluation, urinary organic acid analysis showed markedly increased urinary glycolate (reported as 2000 mmol/mol creatinine, or ~1300 mg/g creatinine) and normal oxalate; blood levels of each were not measured. DNA sequencing also revealed a homozygous splice site disrupting c.814-1G>C variant in *HAO1* shown to abrogate gene function in an in vitro assay. The child was found to have healthy liver function and ultrasound showed normal kidneys, with the only phenotype attributed to the *HAO1* deficiency as the marked glycolic aciduria. However, interpretation of any additional, unknown consequences of glycolate oxidase deficiency in this child was confounded by the co-inherited pLoF in *GMPPA*. Clifford-Mobley *et al.* described a 9 month old girl also born to consanguineous parents with congenital hyperinsulinism due to an *ABCC8* homozygous nonsense mutation^12^. Her evaluation revealed very high urinary glycolate levels, but also with elevated oxalate in spot urines (no blood levels were described), and early stage kidney calcification observed by ultrasound. Importantly, liver glycolate oxidase activity was absent and a homozygous missense variant p.Gly165Cys in *HAO1* was identified. This case was again confounded by co-existing disease, complicating the interpretation of the unexpectedly high urinary oxalate seen given the negative genetic screen for primary hyperoxaluria types 1, 2, or 3 in this infant. Lastly, Craigen reported persistently high urinary glycolate in a 14 month old boy born prematurely who underwent a small intestinal resection for intussusception^13^. Blood glycolate levels again were not measured. The child had normal AGXT activity as measured by liver biopsy and had healthy liver and kidney function. However, no sequencing of *HAO1* or other genes was undertaken to confirm its loss-of-function. Thus, an individual with a confirmed homozygous LoF variant in *HAO1* where phenotyping was not confounded by co-inheritance of additional Mendelian traits remains to be identified.

As part of a large sequencing and health records study in British-Pakistani volunteers^6^, we identified a healthy adult woman with predicted complete knockout of *HAO1* (rs1186715161, ENST00000378789.3:c.997delC, ENSP00000368066.3:p.Leu333SerfsTer4). This variant is predicted to truncate the final 37 amino acids of the protein. Here we describe her medical history, as well as detailed phenotyping and biochemical investigation performed following genotype-guided recall.

## RESULTS

Inspection of blood DNA exome sequencing read data confirmed the homozygous *HAO1* genotype, *Supplementary Figure 1*. Further exome analysis showed the volunteer was 7.4% autozygous at the DNA level, consistent with her parents being first cousins or similar, and the *HAO1* genotype was within an autozygous genomic region. Standard Sanger dideoxy sequencing in a further saliva DNA sample additionally confirmed the genotype, *Supplementary Figure 2.* Neither assay in the different tissues suggested mosaicism. The volunteer was not homozygous for any other rare (minor allele frequency <1%) pLoF genotypes.

Medical history, and National Health Service (NHS) primary care and secondary care electronic health records were reviewed. She was a healthy adult of self-described British-Pakistani ethnicity, in her 5th decade, and a mother with 3 healthy children. An ultrasound of the kidneys, carried out as part of a gynaecology assessment, was normal. She was overweight (BMI, 30-35 kg/m^2^), and other than common non-serious short term illnesses and symptoms associated with pregnancy, had no unexpected medical history.

The cumulative frequency of known pLOF variants in *HAO1* is low: in gnomAD v2.1 (141,456 whole exome/genome sequenced individuals) around 1 in 2700 individuals is heterozygous for a pLoF variant. The rs1186715161 variant was also observed in a single heterozygote in gnomAD. Analysis of pLOF constraint, to assess the degree of natural selection against loss of function, revealed 13 variants observed out of 17.7 expected (obs/exp = 74%, 90%CI 48%-117%), with the confidence interval spanning the range for recessive disease genes (mean obs/exp = 59%) and homozygous pLoF tolerant genes (mean obs/exp = 92%)^10^. The lack of strong constraint makes *HAO1* unlikely to be haploinsufficient, but does not rule out some health effect in homozygotes or a mild phenotype in heterozygous carriers. However, as the expected homozygote frequency in an outbred population would be only ~1 in 30 million individuals, assessing the health effects of loss of HAO1 has been challenging. Importantly, constraint in and of itself does not appear to be associated with likelihood of drug target success^10^ but it is clear that phenotypes associated with variants in the gene encoding a given drug’s target are a good predictor of its adverse side effects^14^. Thus the phenotype of a HAO1 null individual is expected to be informative about the likely safety profile of drugs targeting this enzyme.

Standard clinical venous blood biochemistry including serum sodium, potassium, bicarbonate, chloride, creatinine, ALT, and bilirubin were repeatedly normal at recall and over the previous decade. At recall, the serum anion gap was normal, and plasma and urinary oxalate were both normal. By contrast, and as expected, plasma glycolate (171 nmol/mL) and urinary glycolate (309 mg/g creatinine) levels were markedly elevated (*Table 1*). These results confirm the loss-of-function predicted from genetics, as her plasma glycolate levels were 12 times the upper limit of normal in healthy, reference individuals (14 nmol/mL, n=67) while her urinary glycolate levels were 6 times the upper limit of normal (50 mg/g creatinine).

**Table 1:** Blood and Urine biochemical measurements in a healthy individual with a HAO1 knockout.

| Sample and assay | Level | Reference Range |
| --- | --- | --- |
| *Venous blood* |  |  |
| Plasma oxalate | <1.0 mcmol/L | <1.6 |
| Plasma glycolate | 171 nmol/mL | ≤12 |
| Plasma glycerate | <1 nmol/mL | ≤28 |
| *Urine (single void, same day as blood tests)* |  |  |
| Urine oxalate | 0.16 mmol/L | Not applicable |
| Urine oxalate/creatinine ratio | 20 mg/g | ≤75 |
| Urine glycolate/creatinine ratio | 199 mg/g | ≤50 |
| *24h urine (on a different day)* |  |  |
| Urine oxalate (total/24 hr) | 0.27 mmol/24hr | <0.46 |
| Urine oxalate/creatinine ratio | 23 mg/g | ≤75 |
| Urine glycolate/creatinine ratio | 309 mg/g | ≤50 |

Reference ranges (mean+2sd) derived from studies of >50 healthy adults for each assay at the Mayo Clinic.

Whilst a liver biopsy with direct measurement of enzyme activity was not performed here (because of the risk of biopsy in a healthy volunteer), we can estimate the upper limit of any remaining glycolate oxidase activity from the relationship of *HAO1* silencing to substrate build up (glycolate levels) in rodents and primates, along with levels from *HAO1* silencing in healthy volunteers^2,15^. In a phase 1 trial with the investigational RNAi therapeutic lumasiran targeting *HAO1*, healthy volunteers dosed at the highest dose tested (6 mg/Kg) showed maximal plasma glycolate levels and maximal urinary glycolate levels 3-5 times less than the levels we observed in this individual. Importantly, the 6 mg/Kg dose in the phase 1 study would be expected to silence the HAO1 mRNA at >95% based upon modeling from preclinical results in mice, rats, and monkeys and from direct comparison to clinical data from RNAi therapeutics to other targets that employ the same molecular design and have similar potency^16,17^. Consequently, we estimate that the individual reported here likely retains <2% residual glycolate oxidase activity. Of note, glycolate oxidase is normally found exclusively in the peroxisomes of hepatocytes and requires the C-terminal tripeptide SKI for this targeting^18^. As this subject’s HAO1 protein is predicted to lack the entire C-terminus, it is possible that an error in peroxisomal targeting contributes to this loss of function, in addition to the possibility of nonsense mediated mRNA decay.

## DISCUSSION

The results presented here provide important data to support the safety of *HAO1*-encoded glycolate oxidase inhibition as a potential chronic therapeutic approach for a devastating metabolic disease (primary hyperoxaluria type 1, PH1). Our approach also demonstrates the potential for improved drug discovery through unlocking relevant evidence from the diversity of human genetic variation.

Recent work has both defined the molecular defect in PH1 and provided possible targets to intervention. Multiple approaches are being developed including RNAi therapeutics (e.g. lumasiran, Alnylam Pharmaceuticals; DCR-PHXC, Dicerna Pharmaceuticals) and small molecules (e.g. Structural Genomics Consortium^19^). One promising approach is to inhibit the upstream enzyme, glycolate oxidase (encoded by *HAO1*), to starve substrate for oxalate production and trap it as the more innocuous two-carbon glycolate. Preclinical models have shown the dramatic potential for such an approach for lowering oxalate production by targeting the source of the problem - the defective pathway in liver peroxisomes^2,20,21^ and such an approach has progressed to clinical evaluation (ClinicalTrials.gov NCT02706886, NCT03350451, NCT03681184). Additionally, lumasiran preclinical data and clinical results in healthy volunteers and PH1 patients to date indicate that such an approach is not expected to introduce safety consequences due to *HAO1* silencing itself. However, such results only include chronic studies in non-human species (rodents, monkeys) or relatively short term studies in people, whereas therapy for PH1 is likely to be required lifelong. Consequently, the scientific and medical community have a strong interest in understanding both the potential of *HAO1* silencing for lowering oxalate in PH1 patients along with any long-term consequences of elevations in glycolate levels or other changes in metabolism.

Our report also suggests that any potential other roles for HAO1 in humans beyond glycolate oxidation do not lead to clinical phenotypes, at least in this specific individual with a HAO1 knockout. However samples from this individual might provide an opportunity for further research (e.g. metabolomics/lipdomics of fatty acids) to explore more of human biology.

Any data that informs understanding of the effects of lifelong complete knockdown of a target protein is valuable for the development of a potential therapeutic, both in de-risking the approach and providing potential hypotheses to optimize its development (e.g., biomarkers). Indeed, it is clear that on-target adverse side effects of drugs can often phenocopy the effects of impactful sequence variants in the gene encoding their targets^14^ and therefore the lack of clinical phenotype in this *HAO1* deficient individual is reassuring. Gene silencing based approaches, such as RNAi therapeutics, might especially benefit from such information due to their exquisite specificity. As noted above, though, the chances of finding an individual with a homozygous pLoF mutation in *HAO1* in an unselected outbred population, based upon the heterozygous frequency of such mutations, is approximately 1 in 30 million persons. With no clinically observable phenotype to aid screening, such numbers are unworkable. However, in selected populations where consanguineous unions are more common, the odds of homozygous mutations of all kinds increase profoundly due to autozygosity. However, without specifically looking for an individual with a homozygous pLoF for a specific gene, one is limited to those who present clinically and are indicated for a genetic or metabolic workup. This is seen in interpretation of the previously identified individuals with documented genetic deficiency of HAO1 whose phenotypes were complicated by their co-inherited genetic diseases, as could be expected from consanguineous unions^13^.

Our current report of an HAO1 human knockout whose medical history is not confounded by ascertainment for any phenotype or disease was enabled by a focused study with a unique combination of features. These include i) a population from an ethnic group with high rates of consanguinity such that rare genetic variants occur as homozygotes (due to autozygosity), including homozygous pLoF genotypes (knockouts); ii) excellent e-health record access and analysis (through the NHS); and iii) the ability to directly recall consented subjects for further research by genotype and/or phenotype to established clinical research facilities^22^. This allowed us to find clinically silent, but very informative, variants such as - here - HAO1 deficiency and previously *PRDM9* deficiency^6^. This example shows the potential for, and unique value of, a large international collaborative database of human knockouts to both enable understanding of human physiology and accelerate development of therapies for a broad range of diseases, both rare and common. Ideally, such a collaborative effort would include all, or nearly all, homozygous pLoF variants that are compatible with life but would not require a clinical phenotype for study enrollment. By expanding such a dataset of human knockouts, different genes would be expected to yield different numbers of homozygous knockouts but even an n of 1, as reported here, provides profound insight regarding both its role in human physiology and its potential as a target of intervention. The value of such data cannot be overstated when viewed from its potential to inform, in real time, development of potential therapies for a devastating disease such a PH1. Future work to expand the known universe of human knockouts will certainly provide numerous opportunities to inspire and accelerate disease treatments by leveraging the rich data certain to emerge from focused examination of human genetics and health in such informative populations.

## FUNDING

Genes & Health is funded by Wellcome (WT102627, WT210561), the Medical Research Council (UK) (M009017), Higher Education Funding Council for England Catalyst, Barts Charity (845/1796), Health Data Research UK (for London substantive site), and research delivery support from the NHS National Institute for Health Research Clinical Research Network (North Thames). Born In Bradford is funded by the National Institute for Health Research (NIHR) under its Collaboration for Applied Health Research and Care (CLAHRC) for Yorkshire and Humber, and by Wellcome (WT101597). Genes & Health received funding from Alnylam Pharmaceuticals for work performed in this manuscript. Born In Bradford received funding from Alnylam Pharmaceuticals for data extraction from clinical records. Alnylam Pharmaceuticals directly funded biochemical assays performed at the Mayo Clinic.

Authors from academic or healthcare institutions (Queen Mary University of London, Mayo Clinic, Bradford NHS Teaching Hospitals Foundation Trust, Broad Institute, Kings College London) report no conflict of interest. Several authors are employees of Alnylam Pharmaceuticals as indicated.

We thank Eric Minikel for comments on gene constraint analyses.

## METHODS

The volunteer took part in the Born In Bradford study, and the Genes & Health study. Ethical approval was obtained from Bradford National Research Ethics Committee and the South East London National Research Ethics Committee (14/LO/1240).

Exome sequencing was performed using Agilent in solution capture and Illumina short read sequencing as described in detail in^6^.

Sanger sequencing was performed as described in detail in^6^. Samples were amplified using M13 tagged primer pair below (PCR conditions: Initial denaturation: 10 minutes at 96°C; followed by 35 cycles of 15 seconds at 95°C; 15 seconds at 55°C; 30 seconds at 72°C with a final extension at 72°C for 5mins) before sending for PCR clean up and sequencing with M13F and M13R in-house primers at Source Bioscience. Forward: 5’TGTAAAACGACGGCCAGTTCAAATTCACTTCTCTCCACCA, Reverse: 5’CAGGAAACAGCTATGACCTGGGGCTTAGCTTTCCAGG

Urine and plasma oxalate and glycolate quantification was performed at the Mayo Clinical Labs with samples prepared as instructed for each test (see https://www.mayocliniclabs.com/test-catalog). Urine oxalate (Test ID: ROXU) was measured using a modification of the oxalate oxidase method as previously described^23^. Plasma oxalate (Test ID: POXA) concentrations were determined for an acidified sample (to minimize spontaneous conversion of ascorbate to oxalate) by ion-exchange chromatography using the Dionex ICS 2100 instrument. Urine glycolate (Test ID: HYOX) was determined along with a panel of metabolites (4-hydroxy-2-oxoglutaric, HOG; oxalate, glyoxylate, and glycerate). Briefly, urine samples corresponding to 0.25 mg of creatinine (not to exceed 1 mL of urine) were oximated by reaction with methoxyamine hydrochloride to stabilize one of the target analytes (HOG). The urine was then acidified and extracted with 4:1 ethyl acetate:isopropanol. After evaporation, the dry residue was silylated with 80:20 BSTFA/1%TMCS:pyridine and analyzed by capillary gas chromatography/mass spectrometry (GC/MS) for quantification of each analyte. For plasma glycolate quantification (Test ID: HYOXP), non-acidified plasma samples were spiked with a mixture of stable isotope internal standards and treated with pentafluorobenzyl hydroxylamine HCl to prepare PFB-oxime derivatives of any oxo-acids present. After oximation, available hydroxy groups (alcohols or carboxylic acids) were derivatized with a silylating reagent (N,O-Bis(trimethylsilyl)trifluoroacetamide; BSTFA) to impart necessary volatility and stability for analysis by capillary GC-MS in PCI SIM mode. Quantification was enabled by calibration with use of internal standard, and one quantifier ion and one qualifier ion.

---

**Supplementary Figure 1:**
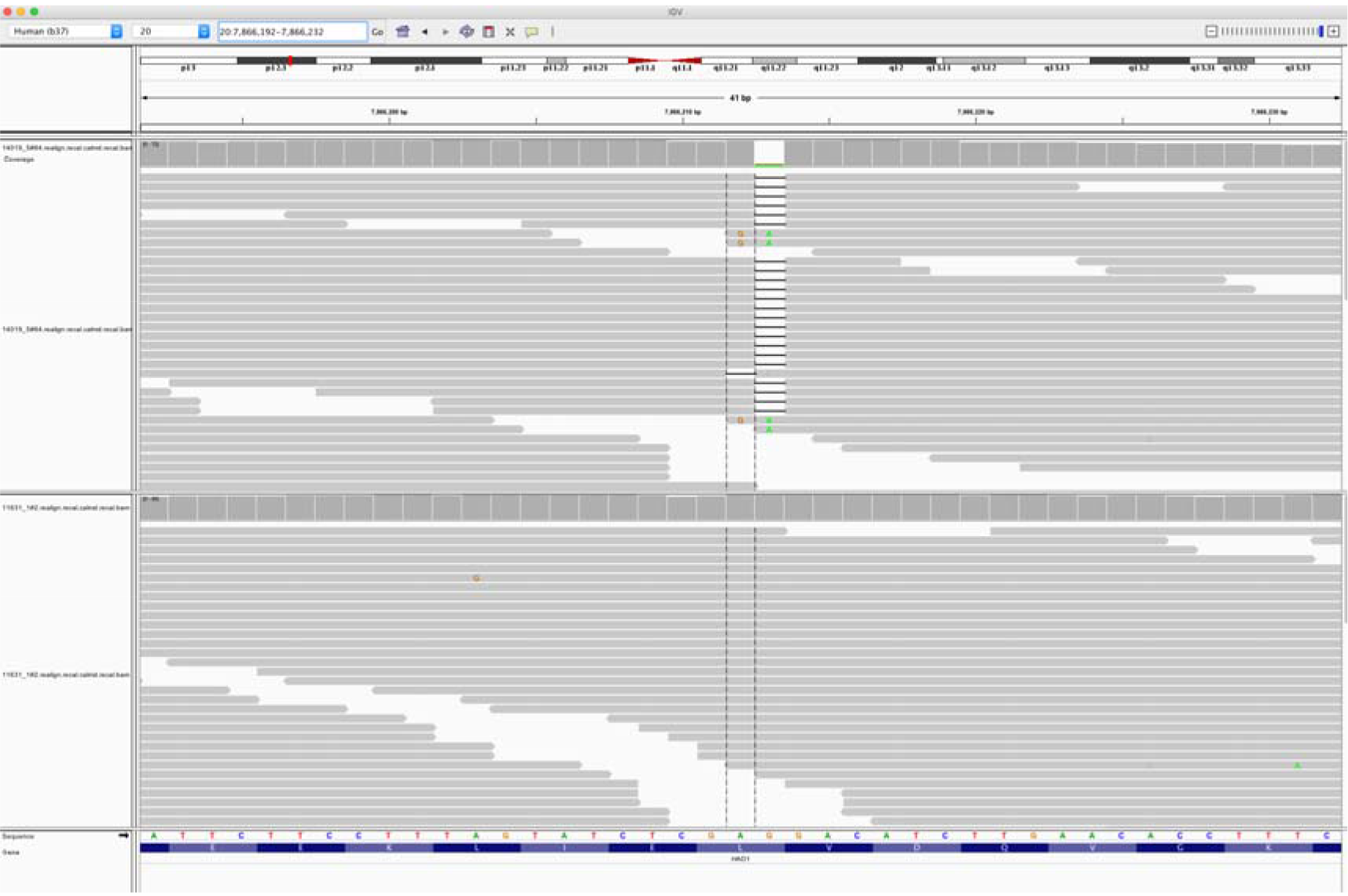
Integrative Genomics Viewer (IGV) image of sequencing alignments around the ENST00000378789.3:c.997delC *HAO1* variant (rs1186715161, GRCh37:20;7866212 AG/A). Exome sequencing performed using Agilent in solution capture and Illumina short read sequencing using blood DNA. Top half of image, individual with homozygous ENST00000378789.3:c.997delC genotype. Bottom half of image, a different individual with homozygous reference sequence genotype.

**Supplementary Figure 2:**
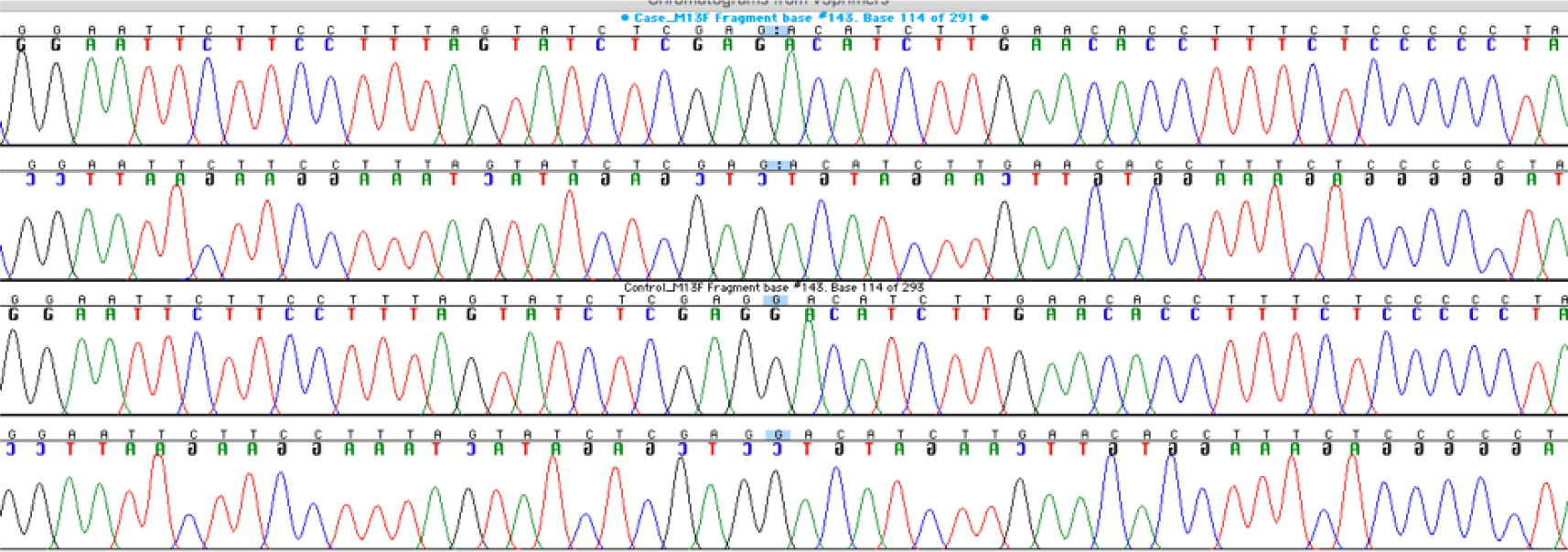
Sanger sequencing around the ENST00000378789.3:c.997delC *HAO1* (rs1186715161) variant using saliva DNA taken at a different timepoint to the blood DNA in Supplementary Figure 1. The top two traces (forward sequencing, then below, reverse sequencing) are of the individual with homozygous ENST00000378789.3:c.997delC genotype. Bottom two traces a different individual with homozygous reference sequence genotype.

